# Kinematics and timing of escape responses in Spotted Ratfish (*Hydrolagus colliei*) and comparison with elasmobranchs and teleosts

**DOI:** 10.64898/2026.04.20.719710

**Authors:** Vincent Mélançon, Heather Bauer Reid, Chelsea Bussey, C. Melman Neill, Jacob L. Johansen, John F. Steffensen, Paolo Domenici

## Abstract

Escape responses are a critical behavioural mechanism influencing survival during predation events. In most species of teleosts and several other lower vertebrates, these responses are triggered by Mauthner cells (M-cells), which generate faster escapes (characterised by higher turning rates and shorter response latencies) than non-M-cell triggered responses. Most adult elasmobranchs lack M-cells and consequently exhibit slower escape response timing than teleosts. Spotted Ratfish (*Hydrolagus colliei*) are a notable exception in that adults possess M-cells, yet their escape response performance has not been explored. Here, we quantify the kinematics and timing of ratfish escape responses elicited by a mechano-acoustic stimulus. We show that ratfish exhibit higher turning rates and shorter response latencies than other adult chondrichthyans, though their response latencies are also significantly longer than those of teleosts. These findings suggest that retention of M-cells confers enhanced escape performance in ratfish, with important implications for their vulnerability to predator attacks.

**Summary statement:** This study reveals that adult Spotted Ratfish (*Hydrolagus colliei*) show fast escape response with a performance that is intermediate between teleosts and previously studied elasmobranchs.

## 1. Introduction

Escape responses, defined as rapid accelerations in response to a threatening stimulus, can vary within and across fish species and provide important information on the vulnerability of fish to predation (Domenici and Hale, 2019; Weihs and Webb, 1984). Escape responses are a type of fast start behaviour, which encompass rapid accelerative movements in a variety of contexts beyond predation (Domenici and Hale, 2019). The study of escape responses incorporates neurophysiology, kinematics, morphology, and behaviour which has led to an extensive body of work related to movement types (e.g., Domenici and Blake, 1997; Domenici and Hale, 2019; Eaton and Hackett, 1984; Eaton et al., 1981; Eaton et al., 2001; Korn and Faber, 2005; Zottoli and Faber, 2000). Escape responses can vary in their kinematics and latency, which largely depend on external factors such as stimulus intensity, distance from the stimulus, and context (e.g., presence of conspecifics; Domenici and Batty, 1997; Domenici and Hale, 2019).

The timing and kinematics of escape responses depends on their underlying neural control, and critically for fish and larval amphibians, on the presence or absence of the Mauthner cells (hereafter M-cells; Eaton et al., 1981; Eaton et al., 2001). M-cells are a paired set of reticulospinal neurons that can be activated by a range of sensory modalities, including mechanosensory, auditory, and visual inputs (Eaton and Hackett, 1984; Eaton et al., 1981; Kohashi and Oda, 2008). Although escape responses can occur in the absence of M-cells, reliance on alternative neural circuits typically results in longer latencies (i.e., slower responses to stimuli) and reduced kinematic performance, as demonstrated in experiments monitoring M-cell activity (Bhattacharyya et al., 2017) or experimentally ablating M-cells (Hecker et al., 2020; Kohashi and Oda, 2008).

The presence of M-cells has been validated in various anamniote species, a vertebrate group comprising fishes and amphibians (Bierman et al., 2009; Eaton et al., 2001; Zottoli, 1978). In chondrichthyan fishes, however, M-cells may be present only during early life stages or absent entirely, as at least seven elasmobranch species appear to lack M-cells as adults (Bone, 1977; Stefanelli, 1980). Only a few species of chondrichthyans have been investigated for the presence of M-cells, however, and escape responses have been quantified in only a small number of elasmobranch species (Domenici and Hale, 2019).

Sharks, rays, and chimaeras are generally large-bodied animals with predatory lifestyles as adults, and thus their escape responses may be understudied in part due to the perception that these species are rarely preyed upon and therefore have limited need for rapid escape behaviour (Estes et al., 2016; Ferreira et al., 2017; Trujillo et al., 2022). However, Domenici et al. (2004) elicited both slow and fast escape responses in adult Pacific Spiny Dogfish (*Squalus suckleyi)*, and Schakmann et al. (2021) subsequently showed that adult *S. suckleyi* exhibit longer escape latencies than teleosts tested at the same temperature. In contrast, Trujillo et al. (2022) reported that neonate tropical reef sharks (*Carcharhinus melanopterus* and *Negaprion acutidens*) can perform fast escape responses with short latencies (5 – 10 ms, comparable to teleosts), which they ascribed to the high experimental temperature (29°C) and the potential presence of M-cells during such early life stages. Collectively, these few studies highlight that escape responses do occur in chondrichthyans and that their timing and kinematics may vary substantially among taxa and life stages. The current paucity of data, particularly for chimaeriforms, is noteworthy given the potential for differences between M-cell mediated escape responses in teleosts and other types of escapes responses in chondrichthyans (Bierman et al., 2009).

Spotted Ratfish (*Hydrolagus colliei*) is a chimaera (Class: Chondrichthyes, subclass: Holocephali) and is among the smaller chondrichthyan species (females can grow up to *ca*. 97 cm; (Tozer and Dagit, 2004). It is a durophagous predator (Huber et al., 2008) distributed along the eastern Pacific coast from Southeast Alaska to the northern Gulf of California (Grinols and Heyamoto, 1965; Wilimovsky, 1954). The depth range occupied by *H. colliei* extends from the subtidal zone (6.0 – 18.5m) down to almost 1000m (Alverson et al., 1964; Cross, 1981; Dean, 1906; Huber et al., 2008; Tozer and Dagit, 2004). Notably, *H. colliei* possesses a robust venomous spine anterior to the first dorsal fin that can inflict wounds and lead to death in small mammals within 48 hours of toxin exposure (Halstead and Bunker, 1952; Tozer and Dagit, 2004). This spine has also been implicated in predator mortality following ingestion, including a documented case in Harbour Seals (*Phoca vitulina*) (Akmajian et al., 2012). Other ratfish predators include elasmobranchs, large teleosts, and some marine mammals (e.g., Pacific Spiny Dogfish [*S. suckleyi*], Soupfin Sharks [*Galeorhinus galeus*], Sixgill Sharks [*Hexanchus griseus*], Pacific Halibut [*Hippoglossus stenolepis*], and Elephant Seals [*Mirounga augustirostris*],) (Andrews and Quinn, 2012; Didier, 1994). Although this venomous spine likely provides an effective defence, venom is generally considered a supplementary rather than exclusive defence strategy (Harris and Jenner, 2019), suggesting that rapid escape responses may also contribute to survival in *H. colliei*.

Importantly, *H. colliei* has been reported to possess M-cells as adults, indicating the potential capacity for fast-start escape responses, although the timing and kinematics of these have not been assessed (Bierman et al., 2009; Zottoli, 1978). Bierman et al. (2009) further suggested that some aspects of the M-cell axon cap in chimaeras may have evolved independently, raising the possibility that M-cell mediated escape responses in holocephalans differs from those of teleosts as well as other chondrichthyans.

The primary goal of this study was to test the hypothesis that escape response timing and kinematics in *H. colliei* resemble those of teleost fishes with M-cells and are faster than those of adult elasmobranchs lacking M-cells. We exposed individuals to a mechano-acoustic stimulus and quantified response latency and turning rates during escape responses. We then compared these values with published data for other fish species tested using comparable experimental approaches.

## 2 Materials and Methods

This research was conducted following guidelines and regulations from the University of Washington Animal Ethics Committee (IACUC permit number 4238-03).

### 2.1 Animal collection and husbandry

Nine (5 females, 4 males) Spotted Ratfish were collected by crane trawls at Point Caution off San Juan Island, Washington, USA (48°33’51.3”N 123°00’50.8”W) from 140 metres depth on 18 July 2023. Fish were held in coolers during transport to Friday Harbor Laboratories (University of Washington), where they were separated by sex and held in two 2.2 m diameter indoor tanks. These tanks were supplied with unfiltered, flow-through seawater (13±1°C). The tanks were housed in a blacked-out room, with lights switched off and opaque coverings lining the windows to reduce stress induced by light exposure. Fish were acclimated to holding tanks for one week prior to experiments and fed live shrimp and clams *ad-libitum*. Fish were fasted for 24 hours prior to the commencement of experiments.

### 2.2 Experimental design

Our design was adapted from Roche et al. (2023) and Schackmann et al. (2021). The experimental set-up consisted of a 2.2 m diameter cylindrical tank with a water level of 30 cm in a blacked-out room. Disturbance to the fish was minimised by surrounding the tank with an opaque suspended tarp. The water level used ensured that a two-dimensional response was the main component of the swimming motion. Escape responses were recorded using a Sony RX100 digital camera at 500 fps mounted 195 cm above the bottom of the tank and near the middle of a 25.4 × 25.4 cm grid covering the tank bottom. This grid allowed for software calibration and appropriate placement during trials. Escape responses were triggered when an individual passed over the grid using a mechanical stimulus (50 mL falcon tube filled with sand) weighing 163.7g. This stimulus was released from a height of 121.5 cm into the water through a tube (tube length: 110 cm, 1.5 cm between tube and water surface, mechanical stimulus stopped at 10 cm below the water surface). Stimuli were manually released when individuals were oriented from within a range of 0 – 90° from the stimulus (i.e., head on or from the side), to minimize variability in stimulus orientation. A mirror was positioned at the water surface near the stimulus tube such that the passage of the stimulus from the tube to the water surface was captured on video. To ensure sufficient lighting, two LED rope lights (120V 60Hz) were attached to the rim of the tank 85 cm from the bottom of the tank and three lamps (home Luminaire brand; 120V 60Hz 3.2W) pointed into the tank at heights between 85 and 100 cm above the bottom of the tank.

Each fish was placed into the trial tank to acclimate for 1 h before experiments began. The acclimation period began with the tank illuminated solely by the LED rope lights. Lamps were turned on 30 – 45 min into the 1 h acclimation period. Following acclimation, the first stimulus was initiated once the individual swam over the stimulus grid area. Subsequent stimulations were provided after a minimum of 10 min delay, if individuals reacted to the first stimulus. After the first instance of a fish not responding to the stimulus, the waiting period between stimuli was increased to 20 min. Trials were run until a fish had a maximum of five successful escape responses or if a fish had three non-responsive trials, at which time trials were terminated. After trials were complete for each fish, each individual was measured for total length and sex was confirmed before being placed in a holding tank. One *H. colliei* individual was also recorded while free swimming undisturbed in order to analyse the kinematics of spontaneous turns as controls (n=6) (Dadda et al., 2010).

### 2.3 Video analysis

Escape responses were recorded individually and analysed using Kinovea (software version 0.9.5). Escape responses were elicited in 29 of 47 stimulation (61.7% escape responsiveness, range per individual 0 – 5 responses). Trials were analysed for individuals with three or more successful escape responses and only when escape responses were fully visible from overhead. All analysed responses were from fish with a minimum of three successful escape responses. Hence one case of two responses in an individual was discarded. An additional response was excluded because the escape response was not clearly visible in the video. Therefore, in total 26 escape responses were analysed across six individuals (3 – 5 responses per individual). Distance was parameterized against the 10 × 10-inch grid on the bottom of the testing tank, and the distance from the stimulus (converted to cm) in each trial was measured as the distance between the stimulus tube and the nearest point of the individual. To determine the latency of each escape response, i.e. the time (in milliseconds) between the moment when the stimulus broke the water surface and the initiation of the escape response, we calculated the time interval between the frame in which the stimulus breached the water surface and the frame with the first detectable movement of the individual following the release of the stimulus. Turning angles were determined by measuring the angle between the line joining the centre of mass (CoM – approximately 0.38 body length from the tip of the individual’s snout using video analysis based on typical values of other fish species Webb, 1978) and the tip of the head, at its starting position and at the end of its turn. The number of frames between the first detectable movement of the individual and when the head stopped moving laterally was taken as the turn duration (in seconds). These values were used to calculate the turning rate (i.e. degrees turned divided by time of the response; degrees/second [deg s^-1^]. The same metrics were measured for control turns.

### 2.4 Analysis

We first compared the experimentally determined turning rates to those of routine turns (controls) using a linear mixed model (LMM) with turning rate as the response variable, type of turn as the explanatory variable (routine turn or escape response), and individual as a random factor. An analysis of variance (ANOVA) was used to determine significant differences (α = 0.05). Model assumptions were verified using diagnostic plots and showed that data did not need any transformation prior to analysis. To compare the obtained turning rates to those of other species, we calculated an expected turning rate value based on the length-turning rate relationship of aquatic vertebrates based on Domenici (2001) using the following formula:(log(*TR_a_*) = −0.8 log(*L*) + 4.3, where TR stands for turning rate and L for length of the animal (cm).

Latency was measured as mean latency (mean of all successful responses from each individual) and minimum latency (minimum value for each individual). Thus, the analysis is based on six data points (one per individual) per latency metric (mean or minimum). We compared our latency values to those of multiple fish species tested in past work carried out at similar temperatures (Table 1). Explicitly, we selected fish species from our study region that had been tested using a mechano-acoustic stimulus and for which minimum latency values were available from Schakmann et al. (2021). We further selected data based on the criteria we used in our own study (i.e., species where each individual was tested multiple times and each individual responded at least 3 times) and extracted mean and minimum individual latencies. When combined, the species we included were the Great Sculpin (*Myoxocephalus polyacanthocephalus)*, Pile Perch (*Phanerodon vacca)*, and Pacific Spiny Dogfish (*S. suckleyi;* Table 1).

**Table 1.** Escape response latencies (mean and minimum ± standard error [milliseconds]) used for species comparison, with testing temperature (°C) and sample sizes (number of individuals and total number of escape responses). Mean and minimum values were calculated after screening the dataset to remove fish that displayed less than three escape responses. Minimum values were taken as the minimum of each individual.

| Species | Mean latency ± SE (ms) | Minimum latency ± SE (ms) | Temperature ($^{\circ}$C) | Sample size (individuals; escape responses) | Source |
| --- | --- | --- | --- | --- | --- |
| Spotted Ratfish<br>*Hydrolagus colliei* | $58.9 \pm 5.9$ | $36.0 \pm 2.7$ | 13.0 | 6; 26 | Current study |
| Great Sculpin<br>*Myoxocephalus polyacanthocephalus* | $44.6 \pm 3.0$ | $20.8 \pm 1.8$ | 13.3 | 7; 31 | Schakmann et al. (2021) |
| Pile Perch<br>*Phanerodon vacca* | $57.9 \pm 5.7$ | $16.7 \pm 4.7$ | 13.3 | 9; 41 | Schakmann et al. (2021) |
| Pacific Spiny Dogfish<br>*Squalus suckleyi* | $100.6 \pm 6.1$ | $66.7 \pm 2.6$ | 13.3 | 8; 33 | Schakmann et al. (2021) |

Escape latencies among species were compared using a linear model (LM) with minimum latency as the response variable and species as the explanatory variable. Significance was determined using ANOVA, and post-hoc tests were performed using the emmeans package (Russel, 2025) with a FDR correction. Model assumptions were verified using diagnostic plots and showed that data did not need any transformation prior to analysis. We then used LMMs to examine the correlation of *H. colliei* (1) latencies and turning rates, (2) distances from the point of entry of the stimulus (distance from stimulus) and turning rates and (3) distances from stimulus and latencies. All models included individual as a random factor. Correlation coefficients (marginal and conditional) were obtained using the MuMin package (Barton, 2024) and slope significance was extracted from the model using the lmerTest package (Bates et al., 2015). When data did not respect model assumptions following diagnostic plot verifications, natural log transformations were applied. All analyses were performed in R version 4.3.2 (R Studio 2024).

## 3. Results and Discussion

In this study, we tested the hypothesis that escape-response timing and kinematics in *H. colliei* resemble those of fishes with M-cells and are faster than those of adult elasmobranchs lacking M-cells. Overall, our results show that escape performance in *H. colliei* is intermediate between that of elasmobranchs and teleosts. *H. colliei* escape responses were characterised by longer minimum latencies than those of teleosts from the same geographical area and substantially shorter than those of adult Pacific Spiny Dogfish (*S. suckleyi*), an elasmobranch that lacks M-cells in adulthood.

Triggered escape responses in *H. colliei* had a mean turning rate of 703.8 deg s^-1^ (275.9 – 1154.1 ± 60.3 deg s^-1^, min-max ± SE) and averaged 41.3 cm total length (35.6 – 48.3 ± 2.1 cm; min-max ± SE), which was significantly higher than those of routine turns (denDF = 31.9, F-value = 59.2, p-value < 0.001; Fig. 1a). Routine turns were approximately 8.5-fold slower than escape responses (82.2 ± 23.0 deg s^-1^, mean ± SE), indicating that the triggered responses represented true escape behaviour (Domenici and Hale, 2019). Turning rates in *H. colliei* were also higher than those reported for adult *S. suckleyi* (*ca* 268 deg s^−1^ and *ca* 471 deg s^−1^ for slow and fast responses, respectively; overall mean *ca*. 364.9 deg s^−1^; calculated based on raw data from Figure 1 in Domenici et al. 2004). The higher turning rates observed in *H. colliei* may be partially attributed to body size differences, as smaller fishes generally achieve higher turning rates (Domenici, 2001; Domenici et al., 2004). Individuals in the present study averaged 41.3 ± 2.1 cm total length, whereas *S. suckleyi* in Domenici et al. (2004) averaged 58.6 ± 4.6 cm (mean ± SE). When evaluated relative to the expected turning rates predicted by the scaling relationship of Domenici (2001), *H. colliei* and *S. suckleyi* achieved 68.7% and 47.5% of expected performance, respectively. Therefore, although both species underperformed compared to broader patterns reported for aquatic vertebrates, *H. colliei* exhibited higher relative agility than *S. suckleyi*.

**Figure 1.**
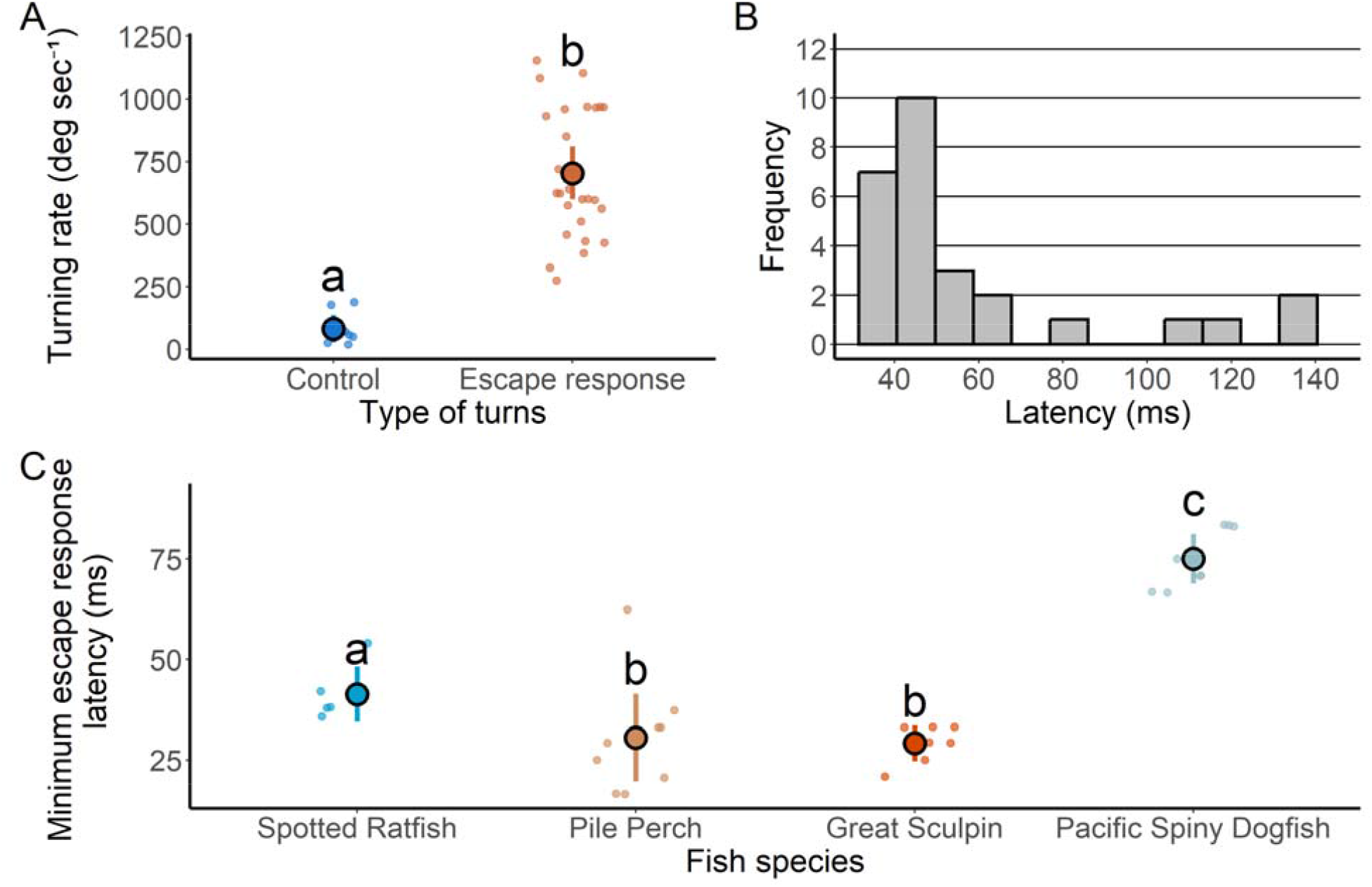
Spotted Ratfish (*Hydrolagus colliei*) escape responses: latencies and turning rates. Panel A) Differences in turning rates (degrees s^-1^) between routine turns (control) and escape responses. Panel B) shows the frequency distribution of latencies (milliseconds) Panel C) Comparison of escape response minimum latencies (milliseconds) among teleosts and chondrichthyans. In both A and C, coloured circles with the black borders represent the mean values while the smaller dots represent the raw data, and vertical bars are 95% confidence intervals. Lowercase letters indicate significant differences.

**Figure 2.**
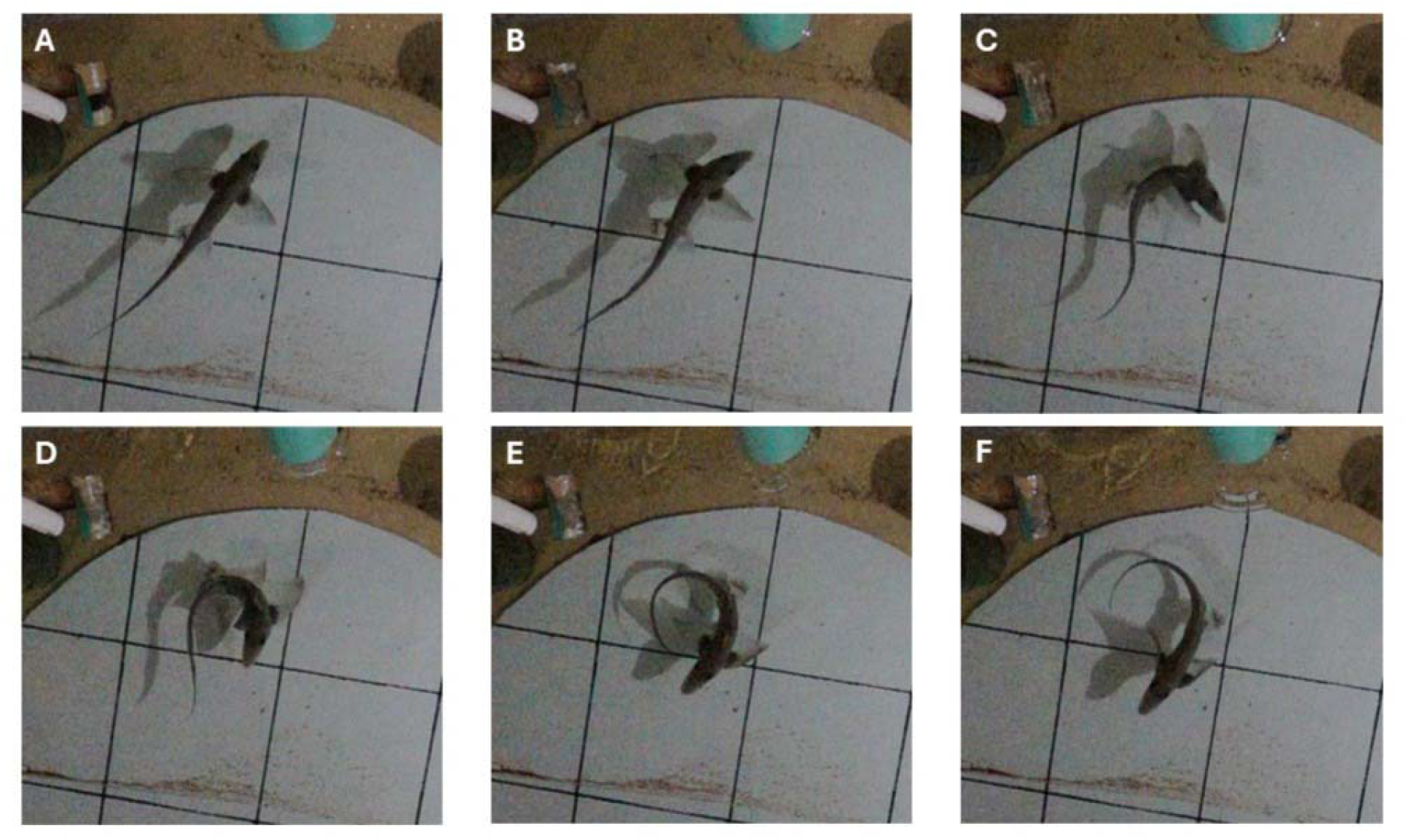
Example of an escape response of a Spotted Ratfish (*Hydrolagus colliei*). The different stages of an escape response following a mechano-acoustic stimulus. From top left to bottom right: A) Moment of impact of the stimulus. The stimulus can be seen in the top left corner though the mirror. The green cylinder is the apparatus through which the stimulus was dropped to prevent individuals from seeing the stimulus as it fell; B) ∼40ms after the first frame; C) 60ms after the first frame; D) 80ms after the first frame; E) 100ms after the first frame; F) 120ms after the first frame. The bottom grid squares are 25.4 × 25.4 cm.

Escape response latencies in *H. colliei* were within the expected range of 5 – 150 ms for escape responses in adult fishes (Domenici, 2010; Domenici and Hale, 2019; Eaton and Hackett, 1984). In our study, mean latency was 58.9 ms (36 – 136 ± 5.9 ms, min-max ± SE), with most values clustering around 40 ms (Fig. 1b). Longer latencies are typically associated with escape responses mediated by alternative neural circuits, while short latencies are more characteristic of M-cell triggered escape responses (Bhattacharyya et al., 2017; Domenici and Hale, 2019). We also found a negative correlation between escape response latencies and turning rates in *H. colliei* (df = 24.7, t-value = −3.11, p-value = 0.005; Fig. S1a), matching the general pattern which shows that shorter latencies tend to be associated with faster turning rates (Bhattacharyya et al., 2017; Domenici and Batty, 1994; Domenici and Batty, 1997; Domenici and Hale, 2019; Hecker et al., 2020).

Minimum latency has been proposed as a more informative measure compared to mean latency because it better reflects maximal escape responses, which is why it was prioritized for species comparison in our study (Schakmann et al., 2021; Trujillo et al., 2022). In *H. colliei*, minimum latency was 36.0 ± 2.7 ms (Table 1), which was slower than the two teleost species examined (Great Sculpin, *M. polyacanthosephalus*, min. latency = 20.8 ± 1.8 ms, p = 0.0436 and Pile Perch, *P. vacca*, min. latency = 16.7 ± 4.7 ms, p = 0.0482), both of which perform M-cell mediated escape responses (Fig. 1c). In contrast, *H. colliei* minimum latencies were significantly shorter than those of adult *S. suckleyi* (p < 0.001; Fig. 1c), which do not possess M-cells.

Comparison with other chondrichthyans also highlights the importance of context, particularly the effect of temperature. A study on two elasmobranch species (Blacktip Reef Shark, *Carcharhinus melanopterus*, and Sicklefin Lemon Shark, *Negaprion acutidens*) found that neonates of these species exhibit much shorter latencies than those that we observed in *H. colliei* (Trujillo et al. 2022). Such short latencies may be partially related to two factors: the potential presence of M-cells in neonate sharks; the temperature at which these neonates were tested (29 °C), whereas *H. colliei* in the present study were tested at 13 °C. Because latency decreases strongly with increasing temperature (Preuss and Faber, 2003; Webb, 1978b), these differences are likely driven largely by thermal effects rather than taxonomic differences alone.

Taken together, our findings suggest that adult *H. colliei* exhibit escape responses with latencies and turning performance that are intermediate between those of M-cell mediated escapes in teleosts, and those of adult elasmobranchs lacking M-cells at comparable temperatures. These results indicate the potential for chimaeras to retain M-cell mediated antipredator responses into adulthood and reinforce the hypothesis that M-cells and associated neuronal circuitry may have evolved independently within holocephalans (Zottoli, 1978; Bierman et al., 2009). The lower escape performance (agility and latency) of *H. colliei* when compared to teleosts may be related to differences in the M-cell axon cap, which is much more developed in teleosts than in chimaeras (Bierman et al. 2009). The retention of rapid escape response capacity into adulthood has implications for the predation risk experienced by *H. colliei*. Predators of ratfish include larger elasmobranchs and teleosts as well as marine mammals (Andrews and Quinn, 2012; Didier, 1994), with their relatively small size likely making them more vulnerable than many other chondrichthyans. In addition, the venom defence of *H. colliei* (delivered through a prominent dorsal spine) requires close proximity to affect predators and therefore is more likely a supplementary or final defence mechanism rather than a primary antipredator method (Harris and Jenner, 2019). With M-cell mediated escape behaviours, *H. colliei* may have greater chances of successfully avoiding predation attempts, which would support the retention of M-cells into adulthood given the relative risks they face.

## 4. Acknowledgements

We dedicate this article to the late John F. Steffensen. We thank all our colleagues during the 2023 Fish Swimming class at Friday Harbor, Washington. We especially thank Mar Pineda and Leon L. Tran. We also thank all personnel of the Friday Harbor Laboratories for their support in fish collection and experiments as well as the use of their facilities and equipment.

## Author Contributions

Vincent Mélançon: conceptualization, data curation, formal analysis, investigation, methodology, project administration, validation, visualization, writing original draft, writing review and editing

Heather Bauer Reid: conceptualization, data curation, preliminary analysis, investigation, methodology, project administration, validation, visualization, writing original draft, writing review and editing

Chelsea Bussey: conceptualization, data curation, preliminary analysis, investigation, methodology, project administration, validation, writing original draft

C. Melman Neill: conceptualization, data curation, preliminary analysis, investigation, methodology, project administration, validation, writing original draft

Jacob L. Johansen: supervision, funding acquisition, resources, manuscript editing John F. Steffensen: supervision, funding acquisition, resources, manuscript editing Paolo Domenici: supervision, funding acquisition, resources, manuscript editing

## Supplementary Materials

**Figure S1.**
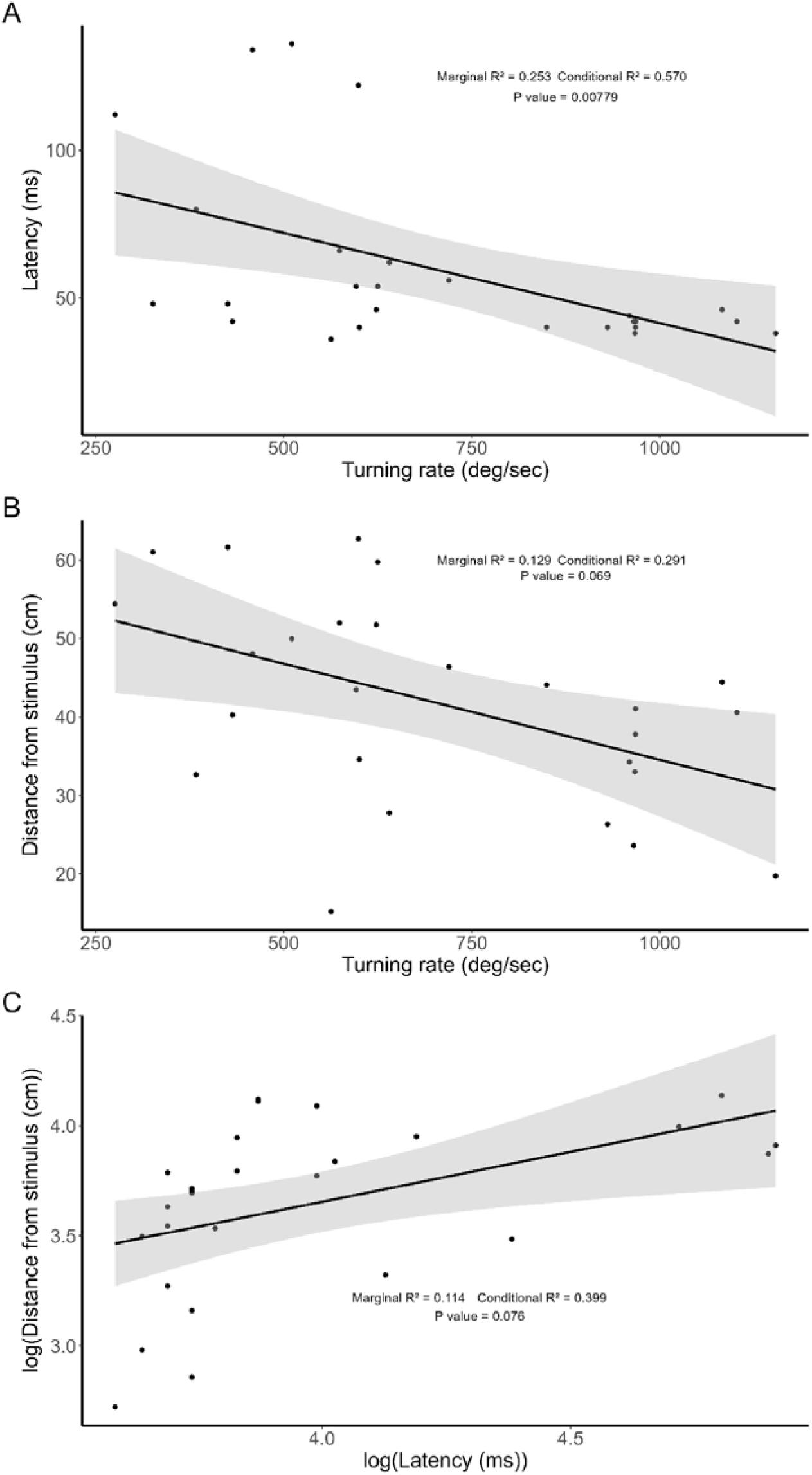
General correlations of Spotted Ratfish escape response metrics. Linear regressions are represented using black lines. Grey area represents the 95% confidence intervals. Panel A) depicts a correlation between latency (milliseconds) and turning rate (deg s^-1^); Panel B) depicts a correlation between the distance from the stimulus (cm) and turning rate (deg s^-1^); Panel C) depicts a correlation between the distance from the stimulus (cm) and latency (milliseconds).

