## Supplementary figures and images for "Kinematics and timing of escape responses in Spotted Ratfish (*Hydrolagus colliei*) and comparison with elasmobranchs and teleosts"

### Fig. S1

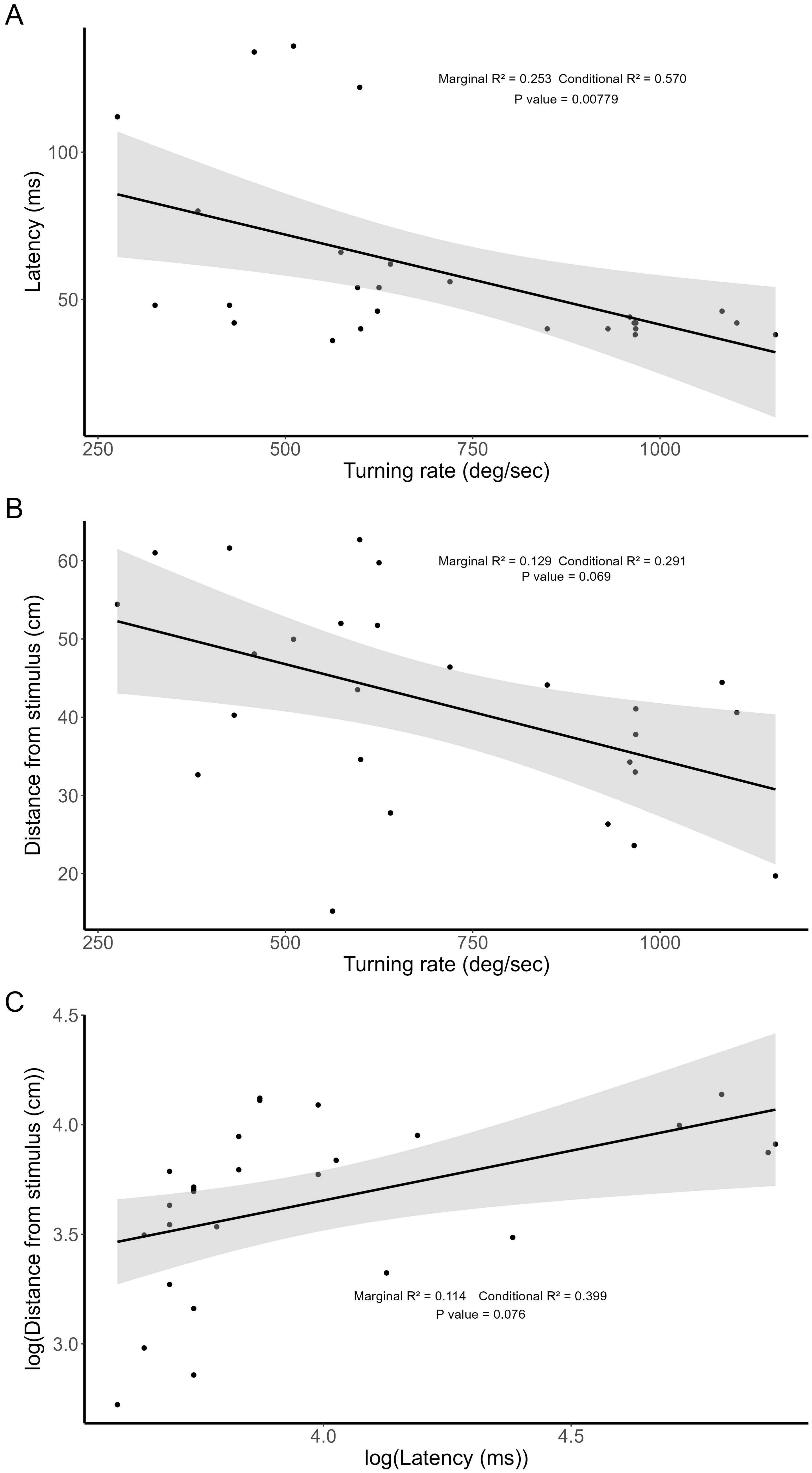
